# A Thermo-responsive collapse system for controlling heterogeneous cell localization, ratio and interaction for three-dimensional solid tumor modeling

**DOI:** 10.1101/2024.12.26.630018

**Authors:** Yu Li, Jordan S. Orange

## Abstract

Cancer immunotherapy using engineered cytotoxic effector cells has demonstrated significant potential. The limited spatial complexity of existing *in vitro* models, however, poses a challenge to mechanistic studies attempting to approve existing approaches of effector cell-mediated cytotoxicity within a three-dimensional, solid tumor-like environment. To gain additional experimental control, we developed an approach for constructing three-dimensional (3D) culture models using smart polymers that form temperature responsive hydrogels. By embedding cells in these hydrogels, we constructed 3D models to organize multiple cell populations at specified ratios on-demand and gently position them by exploiting the hydrogel phase transition. These systems were amenable to imaging at low- and high-resolution to evaluate cell-to-cell interactions, as well as to dissociation to allow for single cell analyses. We have called this approach “thermal collapse of strata” (TheCOS) and demonstrated its use in creating complex cell assemblies on demand in both layers and spheroids. As an application, we utilized TheCOS to evaluate the impact of directionality of degranulation of natural killer (NK) cell lytic granules. Blocking lytic granule convergence and polarization by inhibiting dynein has been shown to induce bystander killing in single cell suspensions. Using TheCOS we showed that lytic granule dispersion induced by dynein inhibition can be sustained in 3D and results in a multi-directional killing including that of non-triggering bystander cells. By imaging TheCOS experiments, we were able to map a “kill zone” associated with multi-directional degranulation in simulated solid tumor environments. TheCOS should allow for the testing of approaches to alter the mechanics of cytotoxicity as well as to generate a wide-array of human tumor microenvironments to assist in the acceleration of tumor immunotherapy.

## Introduction

Novel cancer therapies have advanced tremendously over the last decade led by chimeric antigen receptor (CAR) T-cell therapy (1) and checkpoint inhibitors (2). The efficacy of CAR T cells against solid tumors, however, has generally fallen short of the success seen in preclinical models (3–6). While animal models have always been a mainstay of advancing a cancer therapeutic, a gap in the preceding steps is underscored by limitations of traditional two-dimensional (2D) *in vitro* models, which struggle to replicate the complex environmental conditions inherent in solid tumors (7, 8).

Improvements in the preceding *in vitro* work might lead to animal studies that more effectively translate into successful therapies.

Solid tumors represent a highly specialized 3D environment that includes distinctive biochemical (9, 10), physical (11, 12), tissue (13, 14), and immunological (15, 16) conditions. Recapitulating these characteristics in 2D *in vitro* settings is challenging, necessitating a transition to 3D tumor models, which imposes difficulties in creating multicellular complexity, controlling cell interaction, and maintaining throughput (17, 18). Optimizing these, however, is critical to reduce burdens on animal models and help in ensuring the advancement of the most likely to succeed approaches into animal experiments.

Progress in the development of *in vitro* 3D cell models has been substantial including the use of: 1) microfluidic chips and environmental chambers, which have continually improved in simulating the tumor chemical milieu (19, 20); 2) naturally-derived or synthetic extracellular matrices that have facilitated tissue simulation (21, 22); and 3) cell spheroids and organoids that offer a closely packed cellular environment and some simulated tissue heterogeneity having relevant structural characteristics (23–25).

Importantly, *in vitro* 3D models have been directed as an important accepted alternative and necessary adjunct to *in vivo* animal testing (26).

Despite these advancements and mandates, existing 3D models still face limitations, unable to simultaneously meet all the characteristics needed to allow for experimentally accessible tumor simulation (27). For instance, the preparation of spheroids relies on spontaneous cell adhesion, which limits cell type incorporation and arrangement and fails to mimic or allow for immune cell infiltration (28, 29). Microfluidic devices, though offering active controllability, require highly specialized equipment and are typically quite reductionist (30, 31). This not only limits the scope of experimental measurement but also restricts the throughput of experiments. Organoids and decellularized organs have the potential for intricate cellular and tissue heterogeneity, yet their preparation and maintenance are skill-intensive and experience-demanding, constraining experimental design and throughput (32, 33).

To address these limitations, we have developed a dynamic modeling system that can capture intratumoral cell heterogeneity, while allowing for direct visualization along with precise manipulation of cell positioning, ratios and interactions. Our approach utilized a dynamic scaffold that could mobilize cells via a responsive hydrogel capable of adapting its mechanical properties and internal stress distribution in response to external stimuli. Among various responsive polymers (34), we selected Poly N-isopropyl acrylamide (PNIPAM), which is temperature-sensitive and known for its reversible gelation properties at temperatures above 32°C (35). Specifically, the simplicity of temperature control to promote cell interaction and function along with PNIPAM’s well-documented mechanical properties were appealing (36). We used this tool to build multilayered structures that integrated diverse cellular compositions, mimicking the dense, high interstitial pressure environment encountered within tumors. We also used microbead encapsulation to achieve single-cell-level control of cell distribution and colocalization in an effort to simulate a solid tumor’s complex composition and cellular (micro)landscape (37).

As an illustration, we used these models to explore an accessible behavior of human NK cells that has potential utility in solid tumor cell therapy: multi-directional degranulation. Classically NK cells engage in a cytotoxic process involving a lytic immunological synapse that leads to the mobilization and directed release of lytic granules (38). Those lysosome-related organelles are highly specialized and packed with cytotoxic molecules that enable an NK cell’s destructive capability. After formation of a lytic immunological synapse, the lytic granules converge to the microtubule-organizing center (MTOC) (38, 39) and then polarize towards the triggering target cell to be secreted onto it (40). We and others have shown that this tightly regulated process ensures that the cytotoxic substances are delivered efficiently and precisely, leading to the targeted diseased cell’s rapid destruction without affecting surrounding otherwise potentially healthy bystander cells. Specifically, interfering with this process of convergence results in dispersed lytic granules that are released multi-directionally and can kill both the triggering target cell and neighboring bystander cells (39). While potentially harmful in healthy tissues, multi-directional degranulation could be therapeutically beneficial in the context of solid tumors, for potentially turning immunologically "cold" tumors into "hot" ones, or for eliminating immune-suppressive cells that hinder the anti-tumor response (7, 15). This hypothesis, while suggested in single cell reductionist systems, requires more sophisticated 3D tumor models for effective examination.

Our new *in vitro* model has allowed us to create cell arrangements, visualize them at high resolution, and to begin to evaluate intentionally inducing multi-directional degranulation to enhance NK cell-mediated tumor clearance. While pursued as a use case, it demonstrates the utility of responsive on-demand hydrogel-based 3D model systems and suggests a value of dispersed lytic granules in tumor environments that would not otherwise be appreciable.

## Results

### Responsive hydrogels can be used to embed cells to support the creation of three-dimensional models

A pivotal challenge to modeling solid tumors lies in accurately recapitulating their intricate 3D spatial information, while a substantive challenge to biological studies is derived from needs to accurately visualize and isolate cells in and from 3D models while having control over “time zero”. To create accurate models for biological study, the ability to effectively manipulate relative positions and ratios of diverse cell types within a 3D region is necessary. The ideal models would include additional flexibilities in controlling positioning, differentially labeling, detailed imaging, individual recovery, and on-demand approximation of cells. Thus, we have developed a new strategy using hydrogel matrices that can serve as scaffolds for a 3D model, while allowing the generation of internal stresses that guide cells to desired positions within the model on-demand.

The strategy involves initially encapsulating cells within heterogeneous hydrogel matrices, a portion of which is responsive and capable of rapidly altering its physical properties upon externally controlled temperature change (Fig. 1A). Once embedded, and prior to temperature change, the matrices can then be forged into pre-designed 3D shapes created by extrusion, or emulsification molding. Upon temperature modulation the mechanical properties of the responsive hydrogels change, guiding the cells to specified locations to control their spatial arrangement and generate the “on-demand” contact between cells while thereby precisely controlling “time zero”.

**Figure 1.**
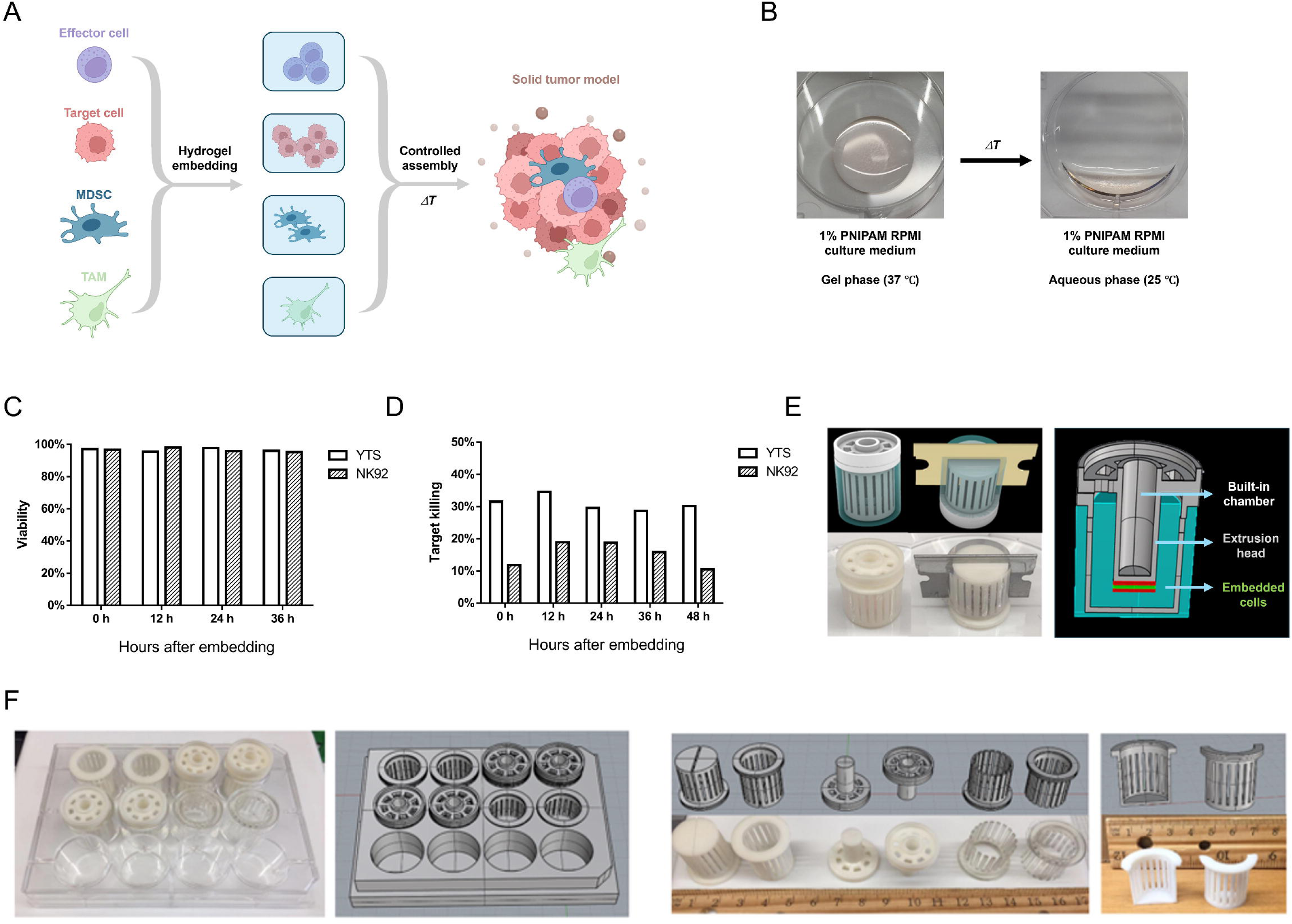
Construction of 3D solid tumor cell models with responsive hydrogel. (A) Our method is based upon individual cell types used to build the 3D solid tumor model being initially embedded within separate responsive hydrogel layers and then combined via a controlled assembly to bring cells into contact on-demand via hydrogel phase transition through external temperature change. This moves the cells via the internal stress within the hydrogel to new positions and contact, thereby enabling control over cell placement and initiation of contact within the 3D model. (B) The hydrogel system was based upon PNIPAM hydrogel that exhibits a gel phase at 37°C and a solution phase at room temperature. (C) The survival and (D) cytolytic function of two commonly used NK cell lines, YTS and NK92, after culture in PNIPAM hydrogel for varying times by uptake of propidium iodide and the specific killing of triggering target cells by ^51^Cr-release assay, respectively. (E) Micromolds with customized cutting slots were designed via CAD and manufactured by 3D printing (left). A built-in chamber was integrated into the micromold to accommodate preheated metal plugs or liquids to facilitate thermal stability for the stacking of cell-containing hydrogel layers (right). (F) Micromold dimensions were scaled to fit into common laboratory consumables, (12-well plates pictured), to promote efficiency and repeatability in 3D cell model experiments. Figure 1A was created with BioRender.com.

Of various responsive polymers to choose from to generate 3D models we selected the temperature-sensitive polymer Poly N-isopropyl acrylamide (PNIPAM). We chose temperature sensitivity owing to the accurate control we have over temperature in our systems and PNIPAM because it is a well-established "smart polymer" having extensive documented mechanical property data (36). For our purposes we took advantage of the fact that PNIPAM undergoes reversible gelation at temperatures equal to or above 32°C when present as a 1% solution in culture medium (Fig 1B).

Although PNIPAM hydrogels have been successfully used for the encapsulation and culture of various cell types (41–43), considering the sensitivity of cytotoxic effector cells to culture environments, we still wanted to validate its biocompatibility with our NK cells. We suspended commonly used NK cell lines in 1% PNIPAM RPMI culture medium and cultured the embedded cells for 24-72h after which the environmental temperature was lowered to facilitate hydrogel dissociation. Thereafter cells were collected and viability and cytotoxic activity against target cells was measured. Flow cytometric analysis after propidium iodide staining demonstrated that the viability of commonly used human NK cell lines (YTS and NK92) was, on average, 96% (range 95∼98%) within 12-36h of PNIPAM embedding (Fig. 1C). Using standard ^51^Cr-release assays the cytotoxic activity of these previously PNIPAM embedded NK cells against target cells was measured and was consistent after 12-36h of incubation and was comparable to the pre-embedded (0h) state (Fig. 1D). Thus, PNIPAM hydrogel exhibited biocompatibility with regards to viability and functionality of NK cells.

To enable precision, standardization, customizability, and reproducibility of 3D model construction, we created standardized micromolds into which PNIPAM cell-containing polymers could be inserted (Fig. 1E. These micromolds, made of photosensitive resin through stereolithography (SLA) 3D printing, presented significant advantages over traditional fused deposition modeling (FDM) techniques. SLA printing occurs simultaneously across the light source screen, in contrast to FDM, which depends on the sequential movement of extrusion nozzles. This parallel action allowed for rapid and uniform production of micromolds. Furthermore, SLA’s precision is dictated by the light source quality and remains consistent across various workpiece sizes, enabling the production of micromolds of different dimensions without a sacrifice in accuracy. Our micromolds have a tolerance as small as +/- 50 microns, eliminating the need for sanding or polishing. To enhance temperature control during the assembly of the model and to help maintain PNIPAM gel state, we integrated built-in chambers into the micromold design that can accommodate preheated metal plugs or liquids to improve thermal stability (Fig. 1E). Additionally, we added standardized positioning pins to enhance interchangeability among different micromolds and to ensure precision during their assembly. We also included customized slots in one of our micromolds so the hydrogel could be sliced to generate a flat surface for placement onto glass to allow for imaging via high-resolution microscopy (Fig. 1E). To provide overall efficiency and promote repeatability, the dimensions of micromolds were scaled for compatibility with common laboratory consumables, such as 24- and 96-well plates (Fig. 1F). This allowed for parallel construction of multiple 3D models using an assembly line-like procedure. Thus, constructing 3D models with responsive hydrogels did not impair cell viability or cytotoxic function and presented options not available in other 3D modeling systems.

### On-demand 3D positioning, function, visualization and isolation of diverse cells via responsive hydrogel generated internal stress and collapse of strata

We next attempted to utilize the compartmentalized responsive hydrogel stresses to determine if we could approximate cells to generate 3D arrangements that could be used to model microenvironments. Thus, we designed a multilayered structure for various types of cells to be embedded in distinct layers of PNIPAM hydrogels. Each layer was produced through extrusion molding using a cyclic micromold, with the thickness of each layer individually controlled by the overall hydrogel volume. The responsive PNIPAM hydrogel layers were encased in a non-responsive hydrogel jacket encapsulating the entire assembly (Fig. 2A). The structure remains stable at temperatures above 32℃, but when dropped below that the PNIPAM hydrogel causes the collapse of the internal layers. Here, to term this process of thermal collapse, in which the internal stresses initially balanced between the two types of hydrogels drive the previously separated cells to aggregate, we called the entire process “thermal collapse of strata, or TheCOS.” The aggregated cells remain within the non-responsive hydrogel jacket and can be sliced for imaging or dissociated for individual cell isolation.

**Figure 2.**
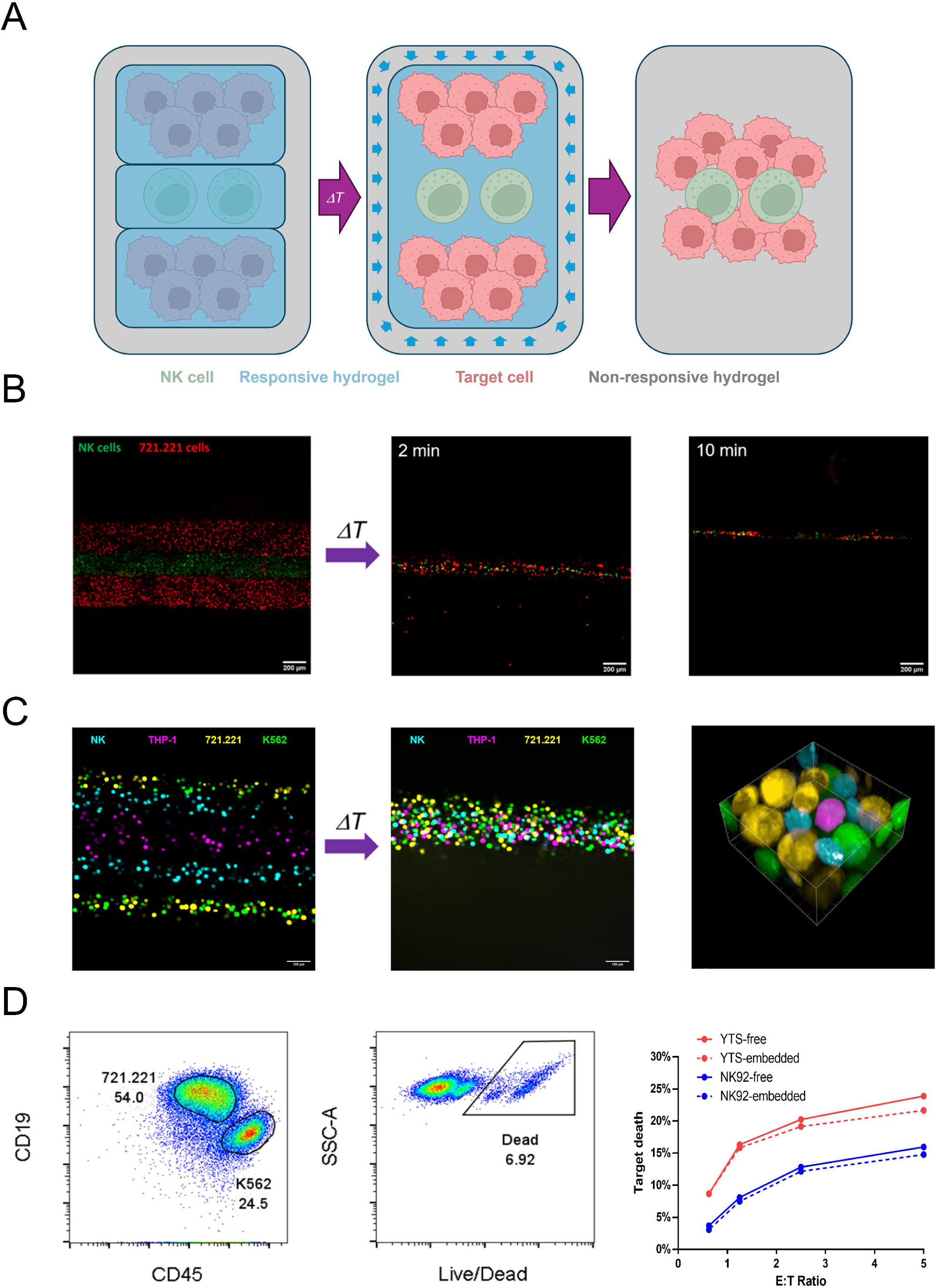
Responsive hydrogel-based strata can enable control over diverse cell positioning, interaction and recollection. (A) Our approach involved embedding labeled cells in layers of responsive within responsive hydrogel scaffolds and forged into multilayer structure which were encased in non-responsive hydrogel. Upon external temperature changes, previously separated cells (example showing a layer of NK cells labeled red between two layers of target cells labeled in green) would be driven by internal stresses to aggregation. (B) Actual microscopy images of a TheCOS stack depicted in the schematic with YTS NK cells labeled in green and 721.221 target cells labeled in green and all of the layers intact at room temperature (left). After the temperature decreased to room temperature the PNIPAM hydrogel collapsed within 10min promoting extensive mixing and mutual contact between the previously separated cells (right). (D) assembly of a 3D models containing five PNIPAM hydrogel layers showing separation at 37 ℃ (left) that collapsed after decrease of temperature to room temperature and demonstrated admixing and contact of the previously separated cells (center). The gel assembly was then imaged at 60x through the z-axis to create a 3-D reconstruction (right). Lytic granules were stained within NK cells by preloading the cells with lysotracker dye and are visualized in white and were in a converged confirmation. (E) Cells were recollected from a TheCOS assembly that had been incubated at 37° for 4 h by digesting the non- responsive hydrogel jacket, washing cells and staining them for flow cytometric analysis. Recollected cells were stained for CD19 and CD45 to allow for identification of the 721.221 and K562 target cells (left) as well as Live/Dead staining to discern those that had been killed (shown for K562, center), allowing for the assembly of killing curves (triggering target cell shown – YTS vs 721.221, and NK92 vs K562) for different TheCOS assemblies having a range of effector to target cell ratios (right, solid lines). Dashed lines demonstrate the same killing assays performed in parallel using admixed cells that had not been embedded. Figure 2A was created with BioRender.com.

As an example and using fluorescently labeled cells (YTS NK cells and 721.221 target cells), we constructed hydrogel assemblies and prepared slides with longitudinal incisions using the cutting slots generated in the micromolds. The sliced hydrogel stack was placed upon a glass slide with the sliced face touching the glass and then visualized via confocal microscopy. Here the PNIPAM hydrogel cell-containing layers (two target cell layers and one NK cell layer) could be visualized within the nonresponsive hydrogel jacket (2B, left). After the collapse of PNIPAM hydrogel via temperature decrease, images revealed extensive mixing and mutual contact between previously separated cells within 2 minutes with completion by 10 minutes (Fig. 2B, middle and right). Thus, differentially labeled cells in ratios determined by concentration in dynamic hydrogels can be brought into contact rapidly on-demand via TheCOS.

TheCOS assemblies can also be modified to utilize multiple layers, including differentially labeled cells at specific densities, in order to simulate a multifaceted tumor microenvironment (TME) that would require many different cell types in differing ratios. As an example, we have chosen to model a 3D environment using 5 layers and four different cell types, each labeled with a different fluorophore in order to demonstrate a simulated TME. We chose a human YTS NK cell, a 721.221 transformed B cell triggering target cell, a malignant K562 erythroleukemia cell (not targeted by the YTS NK cell) and a THP-1 monocytic cell. In this example the two malignant cells were mixed in the outer most top and bottom layers, followed by two internal layers of YTS NK cells surrounding a middle layer of THP-1 cells. As in the previous TheCOS example each cell containing layer was extrusion molded to create an encased stack which was then sliced via the cutting slots and placed at 37° onto a microscope slide and visualized via confocal microscopy. All 5 layers could be seen (Fig. 2C, left), which upon temperature change collapsed (Fig 2C, middle). In this example we also utilized higher resolution and z-axis imaging to visualize the 3D context and create a reconstruction (Fig 2C, right). This allowed for both inter- and intracellular biological visualization. As an example of the former, intercellular contacts could be visualized between an NK cell and each of the other cell types incorporated into the TheCOS simulated TME. As an example of the latter, the lytic granules denoted by lysotracker staining (shown in white) were identified in the foremost NK cell in a converged orientation polarized towards a triggering 721.221 target cell. Thus, TheCOS could be utilized to create complex arrangements of differentially labeled specified cells to generate an on-demand simulated TME.

One advantage of using a hydrogel encasement for TheCOS is that it essentially serves as a container for the PNIPAM layers from which the PNIPAM embedded cells could be liberated after collapse by simply cutting open the stack and dissociating the cells by rinsing with PBS. This would allow for analysis of individual cells using any single cell- or population-based approach. To demonstrate and validate the biological activity of cells in TheCOS, we assembled a three-layer structure consisting of target- NK-target (721.221 target cells and YTS NK cells), collapsed the PNIPAM hydrogel strata by temperature drop and then incubated the assembly for 4h at 37°C. After incubation the assembly was cut through the incision guide, rinsed, and cells collected, and stained for flow cytometric analysis to allow for the detection of each cell type along with viability. Using this approach, the NK cells were easily distinguished from the target cells and both live and dead target cells could be detected (Fig. 2D, left). We also generated different TheCOS assemblies creating effector-to-target cell ratios ranging from 0.5 to 5 and compared these to conventional cytotoxicity assays in which cells were mixed in media. We found that NK cells exhibit cytotoxic activity upon contact with target cells in the embedded state over 4h comparable to that observed in their non-embedded (conventional assayed) state (Fig. 2D, right). With this, the potential of hydrogel-induced stress in manipulating cell positioning and interactions appeared to have no negative effects upon cellular function, providing an iterative basis for further development of TheCOS and related 3D TME models.

### Simulating tumor-infiltrating cells in 3D TME using cell-encapsulating PNIPAM microbeads

Tumor-infiltrating lymphocytes (TILs) in the context of solid tumors have long been pursued and studied and held to be a critical determinant in influencing tumor progression (44). Despite their significance, studies have been challenging owing to a lack of suitable *in vitro* models to reliably provide precise control over the position and interaction of cells in microscale, as well as the timing of interaction. In other words, an inability to place a lymphocyte within a simulated tumor to study its properties. To accomplish this, we prepared dynamic hydrogel microbeads encapsulating single cells to incorporate them into traditional 3D tumor models to control the local distribution of cells and the timing of their contact with the TME.

Using water-in-oil emulsification, aqueous PNIPAM suspended cells were vigorously vortexed in oil to generate microscale droplets suspended in the oil phase. Temperatures gradually rose to 37°, causing the droplets to solidify and form microbeads that encapsulate minimal numbers of cells (Fig 3A). Maintaining the temperature, the microbeads were then centrifugally extracted into the aqueous phase and thoroughly washed (Fig. 3A, right). The size distribution of the resulting microbeads generated using different vortex conditions was evaluated via microscopy (Fig 3B, left). The size of the microbeads was measured via quantitative imaging and the optimal vortex condition for producing microbeads with a diameter of approximately 100 microns was identified (Fig. 3B, right). This specific size was chosen to enhance the likelihood of encapsulating a single cell within each microbead (Fig. 3C).

**Figure 3.**
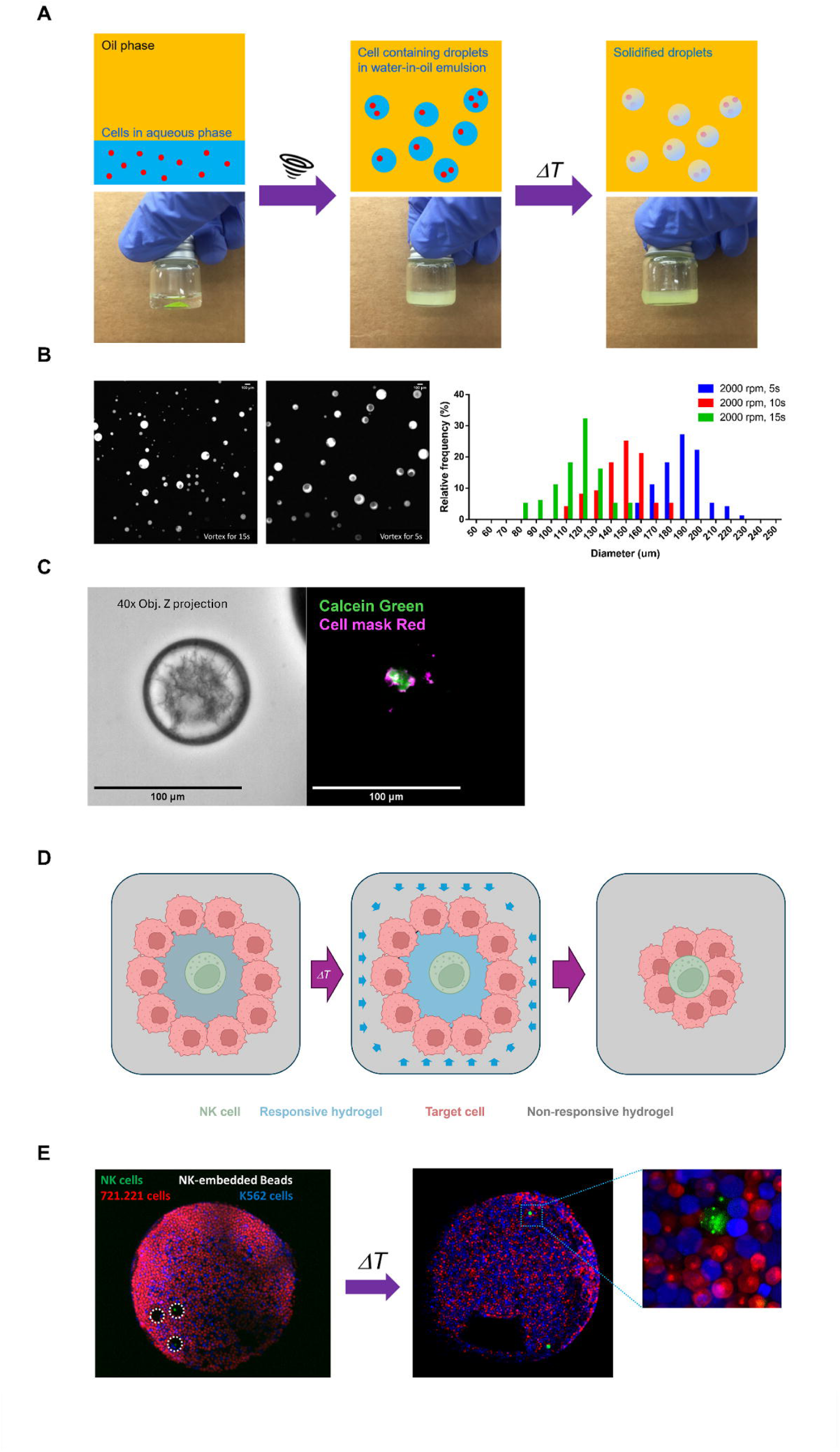
Generation of cell-encapsulating PNIPAM microbeads to simulate tumor- infiltrating cells in spheroids (A) Cell-encapsulating PNIPAM microbeads prepared by suspending cells in aqueous PNIPAM (in this case using hydrogel with a 25° aqueous phase) and overlaying a mineral oil layer (left), which was then vortexed at 25° to emulsify the aqueous cell suspension into droplets (middle), which was then converted into a suspension of hydrogel microbeads in the oil phase by raising temperatures above 37° and converting the aqueous to gel (right). (B) The size distribution of microbeads generated under different vortex conditions assessed by microscopy (left) and measured via image quantitation (right) with 15s giving the most 100µ diameter microbeads. (C) High resolution imaging of microbeads derived from 15s of vortex showing a single viable cellmask red and calcein green-labeled YTS cell contained in a single bead. (D) Illustration of simulated tumor infiltration by an NK cell by introducing an NK cell- containing microbeads into a conventional spheroid. Under typical 37° culture conditions, the microbeads remain in gel phase and encapsulate NK cells (left) until the environmental temperature is lowered to the critical point (middle), leading to collapse of the microbeads and release of NK cells to directly contact target cells within the spheroid (right). (E) Low resolution imaging of NK cell-containing microbeads (outlined in white dashed circles) incorporated into a 721.221 (red) and K526 (blue) cell spheroid at 37° (left). After the collapse of PNIPAM hydrogel via decrease of temperature to room temperature NK cells are found in direct contact with target cells from which they were previously separated (right). Figure 3D was created with BioRender.com.

To simulate a TIL inside a solid tumor, as an example we wanted to try embedding NK cells into microbeads and incorporating them into conventional spheroids. Under typical 37° culture conditions, the microbeads would remain in gel phase and encapsulate the NK cells until the environmental temperature was lowered to the hydrogel’s critical point triggering microbead collapse. After the collapse of the microbeads, NK cells would then be released and come into close contact with target cells within the spheroid (Fig. 3D). Thus, we embedded calcein green-labeled YTS human NK cells in hydrogel microbeads and assembled them into a spheroid consisting of 721.221 susceptible, and K562 resistant target cells that were each labeled with a different vital dye. The incorporation of the NK cell-containing microbeads into the spheroid could be seen by low resolution confocal microscopy (Fig. 3E, left). After the change of temperature and collapse of PNIPAM hydrogel, however, the hydrogel boundary around the NK cell was no longer present and mutual contact could be seen between previously separated NK and tumor cells (Fig. 3E, right).

To model further important elements of the TME via the colocalization of different cells within solid tumors, we extended our microbead strategy by directly encapsulating key low frequency cells for which colocalization would be desired into the same microbeads (Fig. 4A). In order to accomplish this a similar emulsification approach with PNIPAM suspensions was utilized, but for this purpose starting with two different labeled cells that were mixed and vortexed to disperse them into microscale water-in-oil droplets. Once the temperature exceeds the critical point of 37°C, the droplets solidify and encapsulate a mixture of cells with a certain probability. These microbeads would have a high likelihood of containing at least one of each type of labeled cell given the starting ratios used. These microbeads would then be extracted into the aqueous phase and thoroughly washed, after which they could be incorporated into conventional tumor spheroids. Cell contacts between the cells in the microbead with each other and with those of the spheroid, however, would not occur until the temperature was dropped below 32°C. At that point the microbeads would collapse, and the cells within the microbead would be released and come into close contact with each other and the surrounding tumor cells. This would generate an on-demand initiation of a TME including the contact and approximation of lymphocytes and other potentially rare TME cell types to allow for specific and direct biological study.

**Figure 4.**
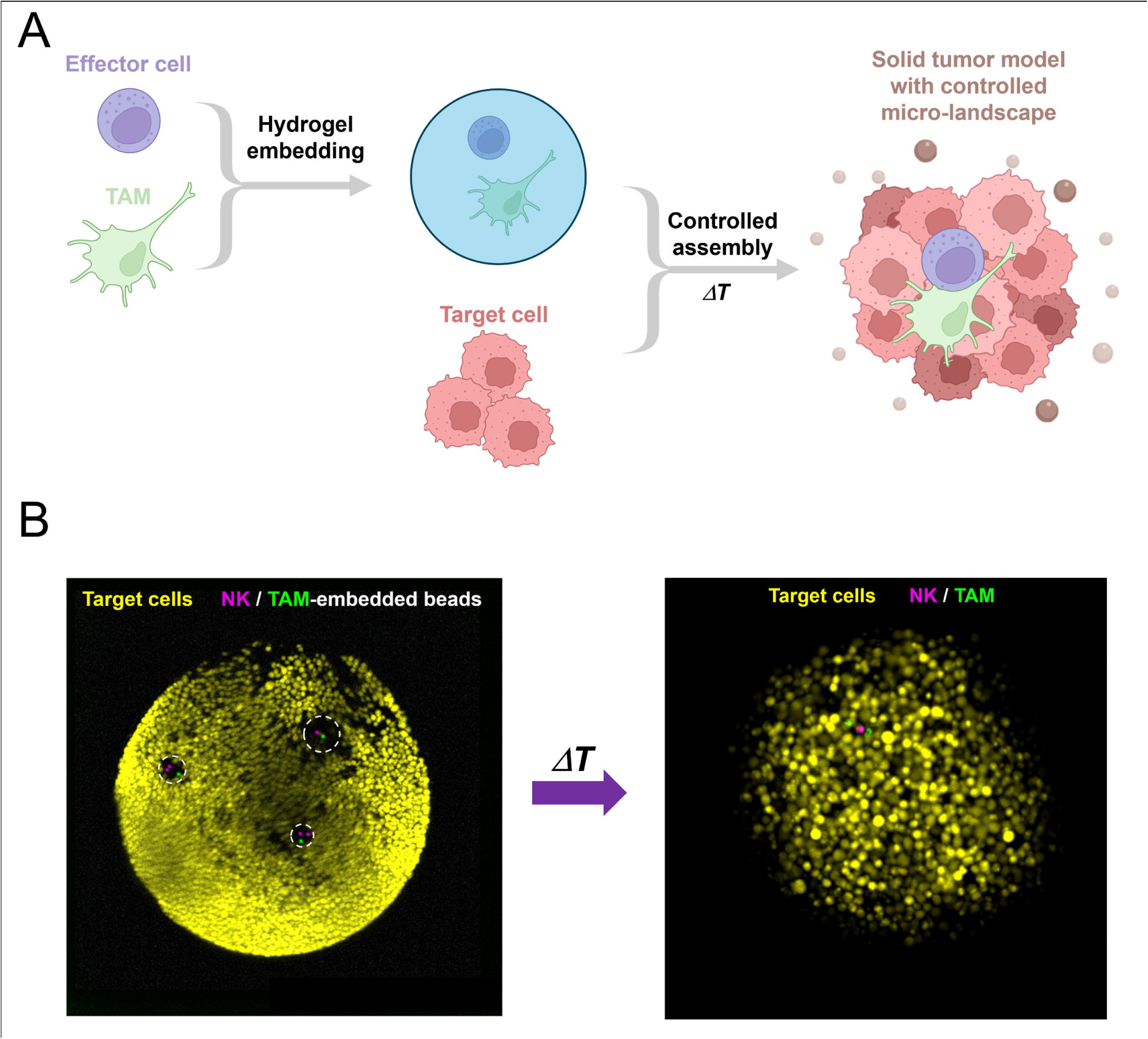
Modeling the colocalization of different cells in solid tumors using cell-encapsulating PNIPAM microbeads (A) Schematic approach for colocalizing an NK cell and TAM in a tumor spheroid in which the cells to be colocalized would be encapsulated in PNIPAM microbeads via the emulsification approach. To promote the embedding of one NK cell and one TAM, equal ratios of the cells would be added to the emulsification procedure. Microbeads would then be incorporated into tumor spheroids after which the temperature would be changed to cause hydrogel collapse and release of the colocalized cells within the spheroid. (B) Formation of a 721.221 target cell (yellow) spheroid with microbeads containing YTS NK (purple) and THP-1 (green) cells outlined in white dashed circles. One microbead contains 1 of each cell and two contain 2 NK cells and 1 THP-1 cell. After the collapse of PNIPAM hydrogel via decrease of temperature to room temperature NK cells are found in direct contact with TAM and target cells from which they were previously separated (right). Figure 4A was created with BioRender.com.

To demonstrate the approach, we created microbeads containing differentially fluorescently labeled YTS human NK cells and THP-1 cells as a surrogate for tumor- associated microphages (TAM). We then constructed a hybrid spheroid model by introducing NK/TAM double embedded microbeads into conventional 721.221tumor cell spheroids. Using low resolution confocal microscopy, we were able to confirm the incorporation of microbeads containing both an NK cell and TAM (Fig. 4B). Importantly the microbead created a zone around NK and TAM that allowed them to avoid contact with each other and the spheroid. After the collapse of PNIPAM hydrogel, mutual contact was formed between previously separated NK cell and TAM and tumor cells (Fig. 4B). Thus, microbeads could be used to precisely manipulate cell positioning on-demand and reproduce intratumoral colocalization patterns amongst typically infrequent cells to allow for direct and kinetic study of their biology from the moment interactions would begin.

### Evaluating the directionality of NK cell cytotoxicity in solid tumor-like environments

In earlier studies, we observed that lytic granules in NK cells undergo convergence to the MTOC and subsequently polarization prior to their degranulation (38). This directs the secretion of cytotoxic cargo into the synaptic cleft, preventing the non-specific killing of innocent bystander cells (39). While we had previously demonstrated this concept in single cell interactions and simple aggregates, it has not been tested in more complex environments. Thus, we hypothesized that bypassing the protective mechanism of convergence and forcing NK cells to degranulate multi-directionally could potentially enhance the destruction of solid tumors and eliminate tumor-resident non-triggering cells (Fig. 5A). This hypothesis, however, has been difficult to examine since conventional assays are unable to simulate the target-rich environment in solid tumors while providing the visualization needed to identify lytic granule positioning. Using, TheCOS, however, we wanted to try and test this hypothesis in solid tumor-like environments and evaluate the possibility of bystander killing by NK cells in a simulated TME.

**Figure 5.**
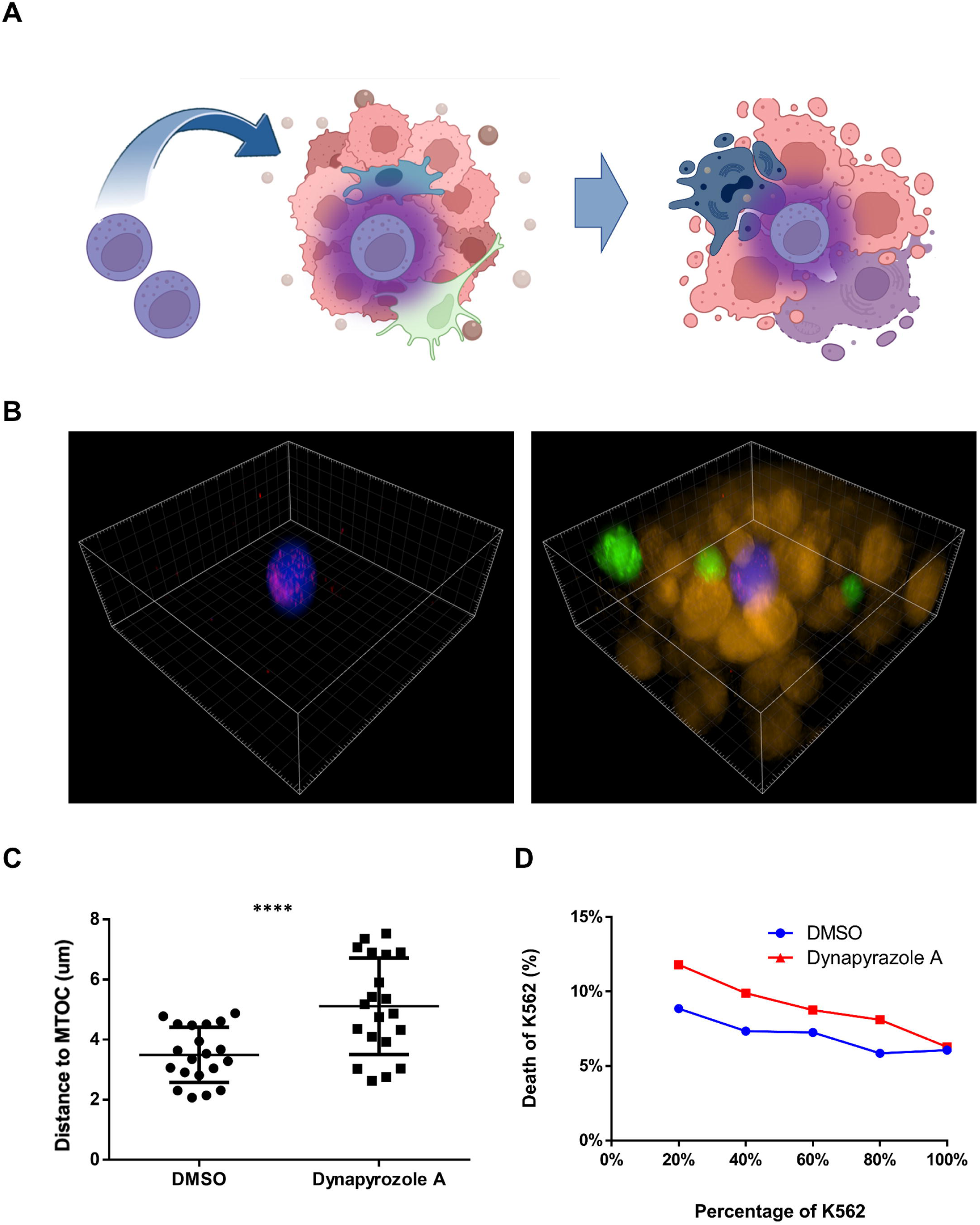
Evaluating NK cell bystander killing in simulated solid tumor environments. (A) Schematic depiction of an NK cell (purple) with its lytic granule forced to degranulate multi-directionally added to a simulated tumor environment of triggering tumor cells (orange) and non-triggering tumor resident cells (green) that could potentially enhance the destruction of solid tumors by eliminating both tumor and tumor-resident non- triggering cells (blebbing and transition to blue). (B) High resolution imaging of a TheCOS experiment with separate PNIPAM hydrogel layers containing YTS (Blue), 721.221 (Green) and K562 (Orange) cells visualized after temperature change and hydrogel layer collapse. In this experiment the YTS cells were pretreated with dynapyrazole A to block convergence of lytic granules (stained with lysotracker, red). Extensive mixing and mutual contact between the different cells (left) as well as dispersion of lytic granules in the YTS cell with all of the others subtracted (right). (C) Mean lytic granule distance to the MTOC of YTS cells, with each point representing the measurement in an individual YTS cell. Dynapyrazole A blocked the convergence of lytic granules in YTS cells compared to DMSO-treated YTS, ****, P < 0.0001, error bars ±SD. (D) The viability of K562 cells in 10 different TheCOS assemblies varying the percentage of 721.221 to K562 cells from 0 to 20, 40, 60 and 80% along with a fixed number of YTS that had been pretreated with either DMSO or Dynapyrazole A after a 4-hour incubation after hydrogel collapse. Death of K562 cells was measured by flow cytometry using Live/Dead dye uptake after dissociation of the TheCOS assembly to create a cell suspension. Bystander killing curves by Dynapyrazole A-treated and DMSO-treated YTS were compared using Mann-Whitney U test that gave a P < 0.001. Figure 5A was created with BioRender.com.

We generated a series of multilayered TheCOS models, encapsulating YTS and target cells at a ratio of 1:20. This configuration aimed to ensure that upon the collapse of the PNIPAM hydrogel, each YTS cell would likely find itself closely encircled by target cells. To modulate lytic granule convergence, YTS cells were pretreated with either DMSO, as a control, or Dynapyrazole A, a dynein inhibitor. Dynein inhibition prevents lytic granules from converging towards the MTOC, leading to their dispersed distribution and non-polarized degranulation that can result in bystander killing (39). 721.221 and K562 target cells were used with the former as the triggering and later bystander target cell since the K562 cells lack the necessary surface activation ligands for YTS cell-induced degranulation. Their proportion varied from 0% to 80% 721.221 cells in separate TheCOS assemblies to allow for different contacts with triggering target cells. In this assembly, after the collapse of the PNIPAM hydrogel, extensive mixing and mutual contact between the YTS and both types of target cells were seen upon high-resolution imaging and 3D reconstruction (Fig. 5B). When YTS cells were used that had been pretreated with dynapyrazole A treatment the lytic granules could be identified in a dispersed configuration (Fig. 5B). Dynapyrazole was effective in blocking convergence compared to DMSO treatment when measured across multiple cells by the distance of the lytic granules from the MTOC in each (Fig 5C).

To evaluate the viability of the target cells TheCOS stacks having the different ratios of 721 and K562 cells were incubated for 4-hours incubation, the hydrogel jacket was digested, cells were harvested and stained for flow cytometry analysis. Viability stains demonstrated that non-triggering K562 cells were largely unaffected by control-treated YTS cells and even in up to 80% presence of 721.221 triggering target cells.

When the YTS cells were pretreated with dynapyrazole A, however, there was a significant increase in K562 cell death in the increasing presence of triggering 721.221 target cells (Fig. 5D). Thus, YTS cells with dispersed lytic granules when triggered in a TheCOS simulated 3D TME mediated bystander cell killing in a dose-responsive manner.

Compared to the classical unidirectional degranulation process resulting from polarized lytic granules, the molecular details and mechanisms of multi-directional degranulation remain unknown. For instance, is multi-directional degranulation more akin to the secretion process observed in mast cells, or is it merely the ’leaking’ of a few granules outside the synaptic region? It has been difficult to address this question using reductionist aggregations of single cells in 2D layers. That said, clarifying this question is crucial and especially in 3D if there are going to be considerations given to using this strategy in trying to optimize cytotoxic cell therapy for solid tumors.

To map the directionality of the degranulation of dispersed lytic granules, we conducted live cell microscopy within our TheCOS model using YTS cells treated with dynapyrazole and triggered by 721.221 target cells and in the presence of K562 bystander cells. In this case the target cells were membrane dye-labeled to allow for their identification and also calcien-loaded to allow for the determination of their viability. When a labeled cell lost calcein it was considered a death event. The death of bystander K562 cells by a contacting YTS cell that was also in contact with a 721.221 target cell was considered a multi-directional degranulation. In a TheCOS stack, Dynapyrazole-treated YTS cells could be visualized contacting both 721.221 and K562 and the loss of the Calcein from both could also be identified denoting both triggering target and bystander cell killing (Fig. 6A). Anchoring on NK cells, we recorded the locations of neighboring dead target cells and designated the position of a dead 721.221 cell as the orientation of the lytic immune synapse. The relative directions of dead K562 cells to this reference were then measured as the angle of degranulation relative to the lytic synapse (Fig. 6B). In Dynapyrazole treated YTS cells the angular distribution of bystander killing was present in all angles from the lytic synapse (Fig 6C) but was not completely random as there was a bias in the direction of the synapse (Fig 6D). In other words, NK cells with dispersed lytic granules demonstrated bystander killing in all directions with some increased killing of non-targeted cells that were closer to the triggering target. This also distinguishes dispersed lytic granule degranulation from the random degranulation that would be expected in an unrestrained multi- directional release (such as in a mast cell) and suggests a unique mechanism underlying the multi-directional degranulation of NK cells.

**Figure 6.**
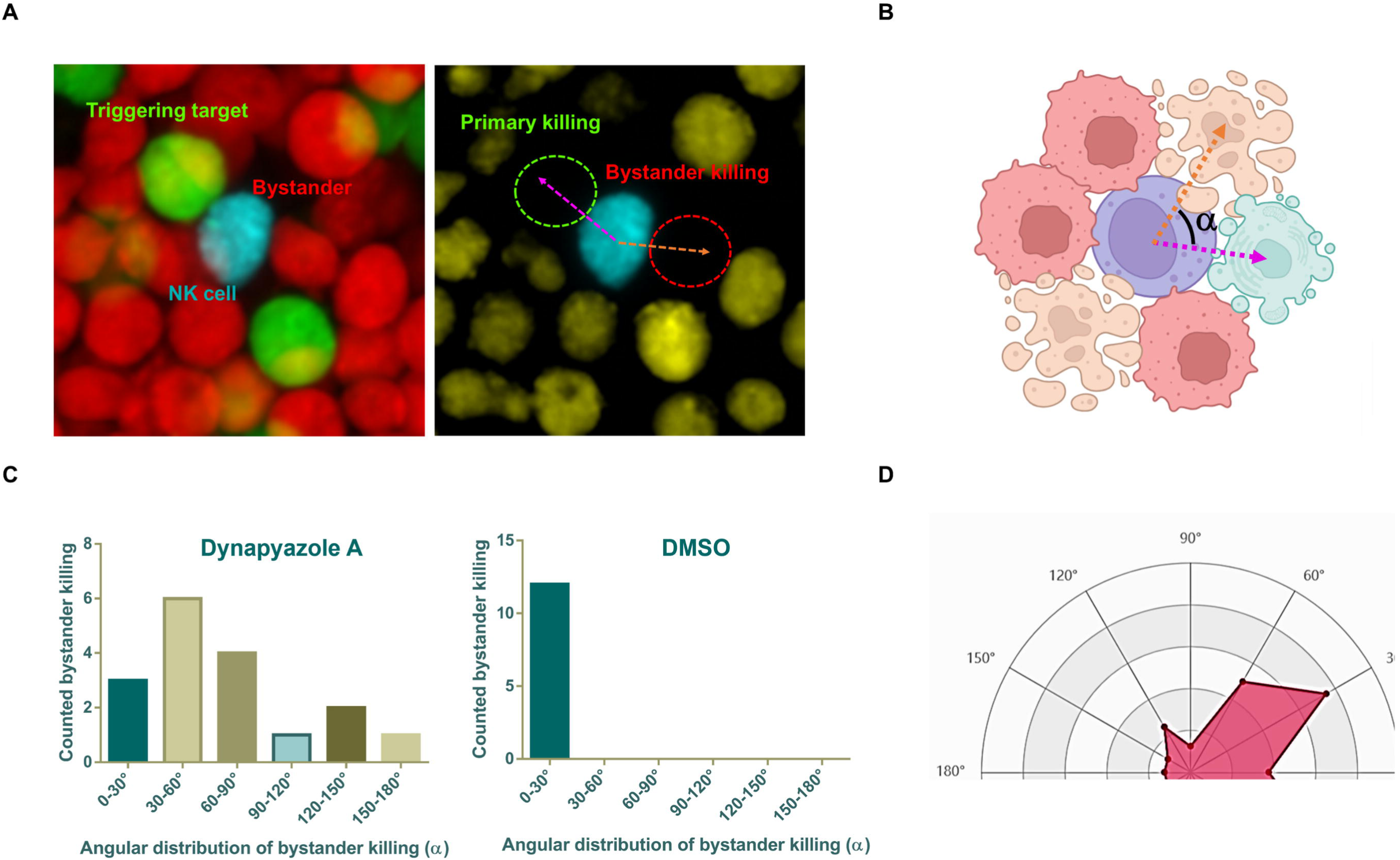
Directionality mapping of bystander killing in modeled solid tumor-like environments (A) To map the directionality of bystander killing in a simulated tumor environment mediated by NK cells having blocked lytic granule convergence a TheCOS assembly was created using Dynapyrazole-treated lipid-dye labeled YTS (Cyan) in separate layers from 721.221 (Green), and K562 (Red) cells that were collapsed, incubated for 4h and imaged (left). The target cells were also labeled with cytosolic dye (orange) and dead target cells identified by those that lost cytosolic dye (right), with the example of the killing of a triggering target cell (green circle) and bystander cell (red circle). (B) The directionality of bystander killing was quantified as the relative directions of dead K562 cells to the orientation of the primary immune synapse formed with the triggering 721.221 cell. (C) Frequencies of different angular directionality of bystander killing events by Dynapyrazole-treated YTS cells (left) or control-treated (right) YTS cells categorized into 6 directional categories, (D) shown also as a radar chart. Figure 6B was created with BioRender.com.

## Discussion

The development of meaningful 3D tumor models is pivotal for the advancement of both basic cancer biology and clinical investigation (45, 46). These models offer a more accurate replication of the complex architecture and heterogeneity of tumors than traditional 2D cell cultures. In pre-clinical contexts, 3D tumor models present value for drug screening and the pursuit of personalized medicine approaches (47, 48). In particular, they can support the evaluation of therapeutic agents in environments that more closely mirror actual tumors to enhancing the predictability of drug efficacy while potentially reducing some of the exploratory burden upon animal models (49). With regards to basic cancer biology, advanced 3D models provide deeper insights, including cell-cell interactions (50), dynamic crosstalk between tumor associated cells (51), and mechanisms of progression and metastasis (52). Capturing interactions within the TME is vital to strategizing improved therapies and understanding how the microenvironment influences cancer progression and treatment efficacy.

Despite their benefits, current 3D tumor models have numerous challenges (53). Firstly, replicating the complexity of TME, especially the role of the immune system in cancer progression and treatment response. Secondly, standardizing models for reproducible application in different research laboratories and high-throughput screening. Thirdly, the considerable resources and specialized expertise in preparation and upkeep of cutting-edge 3D models. Finally, a general inability for models to be precisely controlled in time while enabling direct visualization at high resolution.

Therefore, an ideal 3D tumor model should not only accurately replicate the complex tumor microenvironment, including interactions among immune and stromal cells, but also be scalable for high-throughput applications, ensure reproducibility, and incorporate advanced imaging technologies. Equally important is the balance between complexity and practicality, making these models accessible for both fundamental research and the pre-clinical development of therapeutic strategies.

In our work, we utilized hydrogel matrices as scaffolds to create 3D multicellular structures and achieved dynamic spatial control of cells by leveraging their internal stresses. In our design, PNIPAM hydrogels were used to embed distinct cell types individually, turning them into building blocks for 3D assembly. Since these hydrogels can undergoing a gel-to-solution phase transition in response to temperature change (36), it allows for a spatial collapse subjecting cells to a rapid yet mechanically gentle relocation. We also designed and manufactured a series of molding tools to ensure the precision, speed, and repeatability of 3D model construction enabling a broad-scale mixture of various cells at specified ratios, while simulating the closely packed and high interstitial pressure environment faced by cells in solid tumors. Finally, our system also allows for the easy isolation of cells from the matrix for population-based or single cell analyses thus combining some of the utility 2D reductionist systems with the special benefits of 3D modeling.

Theoretically, stimuli-responsive (54–56) and light-responsive hydrogels (57–59) could produce similar effects. However, these systems typically depend on the action of enzymes or other reactive chemicals to break down the hydrogel backbones, or they utilize spatially modulated UV light to degrade hydrogels at specific sites. As a result, stimuli-responsive hydrogels are inherently slow due to the rate-limiting nature of depolymerization reactions. Efforts to accelerate this process by increasing the concentration of enzymes or reactive chemicals are often counterproductive, causing higher costs and increased toxicity (34). Similarly, the degradation rate of UV-responsive hydrogels is directly tied to the intensity of UV light, which is constrained by UV-induced phototoxicity (60). As an alternative strategy, microfluidics systems utilize carefully designed microchannels and chambers to manipulate cell positioning and contact (61). Though cells can be moved rapidly and with minimal mechanical stress through controlled flows, these devices, which are often chip-based are typically highly customized making them expensive and inflexible. In contrast, our approach relies solely on ambient temperature control offering several advantages: 1) ultra-low toxicity; 2) minimal equipment; 3) cost-effectiveness; and 4) compatibility with common imaging platforms; thus offering a practical solution for dynamic 3D tumor modeling.

In the present study, as a use case and application we used our hydrogel system to evaluate a particular characteristic of cytotoxic effector cells. During target cell-killing they characteristically converge their lytic granules towards the MTOC. Propelled by dynein motors, the lytic granules migrate along microtubules toward the minus end, to reach the MTOC (38). This behavior of cytotoxic effector cells allows them to precisely eliminate the targeted diseased cells while sparing healthy, neighboring cells (39). This precision minimizes collateral damage to surrounding tissues and provides an efficiency to cytotoxicity and presumably superior function in surveillance. Disruption of this convergence process leads to a multi-directional degranulation of cytotoxic cells, increasing the unintended destruction of adjacent healthy cells (39). In certain scenarios, however, a more multi-directional mode cell killing may be desirable and therapeutically beneficial. In particular, in the context of solid tumors, tumor-infiltrating lymphocytes often encounter a suppressive microenvironment that hampers their activation and degranulation capabilities (15, 51). Provoking multi-directional degranulation within such a hostile environment may unleash suppressed cytotoxicity, maximizing the total impact of each round of degranulation and killing additional tumor and other TME-resident cells through increased collateral damage.

In this study, we treated NK cells with dynapyrazole A (62), a dynein inhibitor that disrupts the convergence of lytic granules to the MTOC, and evaluated them in a simulated TME created by TheCOS. Therein, we observed NK cells undergoing multi- directional degranulation outside the immune synapse, inflicting damage on neighboring bystander cells. This suggests the potential of deliberately inducing multi-directional degranulation to boost NK cell-mediated tumor clearance after NK cells have entered a solid tumor and been specifically triggered. This approach serves as another avenue for amplifying cytotoxicity by circumventing an intrinsic protective mechanism and holds promise for refining current therapies.

The use of TheCOS modeling in this context has also provided some additional insight into the function of the lytic immunological synapse. Traditionally, it was believed that degranulation occurred centrally within the lytic synapse, close to where ligand-bound activating receptors were (63), and in an area characterized by relatively low levels of cortical filamentous actin (F-actin) that would provide lytic granules plasma membrane access (64). We found that multi-directional degranulation could occur, but was not entirely random favoring the direction of the immune synapse, despite this area having higher levels F-actin (40). This preference could suggest that F-actin may play a facilitating role in degranulation rather than merely serving as a barrier to the granule approach. For example, it has been shown that integrins, which are directly linked to the F-actin cortex, mechanically license membrane subdomains for degranulation (65).

Specifically, the integrin-mediated mechanical coupling between the cytotoxic cell and the target cell allows the former to "feel" the presence of the target by pushing or pulling against it. This may explain the observed discrepancy between the biased directionality of degranulation and the dispersed distribution of lytic granules, highlighting the critical role of 3D tumor models in advancing insights into the basic immunooncology.

We could conceive many use cases for TheCOS and ones that expand far beyond immunology and cancer biology. The ability to layer cells upon demand and create complex arrangements with specified ratios could be helpful to constructing synthetic tissue, while allowing for the experimental observation, and visualization, of the process in order to provide for iterative improvement. The precise control of the initiation of cell approximation and ratios of cells, while allowing for direct visualization and single cell isolation can have broad applications that we hope will be useful to cell biology and efforts in clinical translation. We also hope that it can represent utility for the rapidly evolving field of 3D cell modeling and allow for greater effectiveness when pre-clinical experiments turn to the use of *in vivo* animal models.

## Supporting information

Key Resources Table

## Acknowledgments

The authors wish to acknowledge Dr. Luis A. Pedroza, Frederique M. van den Haak for valuable discussions and comments on the experiments and manuscript. This work was supported by the National Institutes of Health (NIH R37AI067946) to JSO.

## Materials and Methods

### Cell Lines

The NK cell line YTS (RRID: CVCL_D324), NK92 (RRID: CVCL_2142) and the target cell lines 721.221 (RRID: CVCL_6263), K562 (RRID: CVCL_K562), and THP-1 (RRID: CVCL_0006) were obtained from ATCC or a collaborating lab. These cell lines were validated by phenotypic markers and routinely tested for mycoplasma contamination. YTS, 721.221, K562, and THP-1 cells were cultured in complete RPMI-1640 medium (Gibco) supplemented with 10% fetal bovine serum (FBS) (Gibco) and maintained at 37°C in a humidified atmosphere with 5% CO₂. The NK92 cells were cultured in alpha-MEM medium (Gibco) supplemented with 12.5% FBS, 12.5% horse serum (Gibco), 0.2 mM inositol, 0.02 mM folic acid, 0.1 mM 2-mercaptoethanol (Sigma), and 2 mM L-glutamine. Additionally, 100 U/mL recombinant human IL-2 (PeproTech) was added to maintain cell activity and proliferation.

### Design and Fabrication of Micromolds

Micromolds for 3D tumor models were custom-designed using Rhinoceros 6.0 software (Robert McNeel & Associates) to create digital models, which were exported as standard STL files. Design schematics and CAD files are provided in the supplementary materials. The STL files were imported into a Creality LD-002R SLA 3D printer (Shenzhen Creality 3D Technology Co.), and micromolds were fabricated using ELEGOO Standard UV-Curing Resin (ELEGOO Inc.). After printing, micromolds were washed in isopropyl alcohol (Sigma-Aldrich) then subjected to 405 nm UV light exposure in an ELEGOO Mercury Plus curing station (ELEGOO Inc.) for post-processing. Accuracy and tolerance (<0.1 mm) were assessed using a digital caliper (Mitutoyo) and verified against the design specifications. This ensured the micromolds met the precision requirements for 3D tumor model formation.

### Preparation of 3D Cell Strata

To prepare 3D cell strata, 1% agarose gel solution (Sigma-Aldrich) was poured into a pre-assembled micromold and allowed to solidify at room temperature (∼25°C) for 20 minutes. Once solidified, the upper micromold was gently removed, exposing the agarose gel base (lower die). Cell suspensions were prepared by resuspending cells in 1% PNIPAM complete RPMI-1640 culture medium (described previously) at a temperature slightly below the phase transition temperature of PNIPAM (∼32°C). The exact volume of cell suspension required for each layer was calculated based on the desired thickness. The cell suspension was pipetted into the lower die, and the upper micromold was carefully aligned and pressed down until it made contact with the agarose gel base. The assembled mold was transferred onto an isothermal plate (37°C, Thermo Fisher Scientific) and incubated for 5 minutes to induce PNIPAM gelation. After gelation, the upper micromold was gently removed to reveal the newly formed cell layer. This process was repeated to construct additional layers, achieving the stratified structure. To seal the cell strata, 1% agarose solution (∼45°C) was poured into the lower die, covering the cell layers. The assembly was cooled on the isothermal plate for 20 minutes to allow the agarose to solidify. The final gel assembly (agarose + PNIPAM) was demolded using a specialized demolding tool, while maintaining contact with the isothermal plate (37°C) to preserve the structural integrity of the layers. To collapse the cell strata, the assembly was cooled below the PNIPAM transition temperature (≤32°C) for 5–10 minutes. For incubation, the gel assembly was transferred to an incubator (37°C, 5% CO₂) for 4∼12 hours.

### Preparation of Microbeads and Spheroid Models

Spheroid models were generated by first suspending cells in a 1% PNIPAM solution (Sigma-Aldrich) at a concentration of [specify cell density, e.g., 10 cells/mL]. Both the cell suspension and mineral oil (Sigma-Aldrich) were pre-warmed to 28°C on an isothermal plate (Thermo Fisher Scientific). To create emulsified droplets, 100 µL of the cell suspension was added to 2 mL of mineral oil in a 15 mL conical tube (Corning) and vortexed at maximum speed using a vortex mixer (VWR) for 30 seconds. The emulsion was incubated in a 37°C water bath for 3 minutes with gentle shaking (80 rpm). The vessel was dried externally and placed on an isothermal plate (40°C) for PNIPAM gelation. Gelled microbeads were separated from the oil phase by transferring the emulsion into pre-warmed PBS (37°C, Gibco) using a micropipette. Serial dilutions were performed in 96-well low-adhesion plates (Corning) to isolate individual microbeads. All steps were conducted on an isothermal plate (40°C) to maintain consistent temperature conditions. Isolated microbeads were co-cultured with target cells in a low-adhesion 96- well plate (Corning) for 72 hours in an incubator (37°C, 5% CO₂) to promote spheroid formation. To encapsulate spheroids, 2% agarose hydrogel (prepared at 45°C, Sigma-Aldrich) was added to each well in equal volume. The plate was cooled on an isothermal plate (40°C) for 30 minutes to solidify the agarose. The embedded spheroids were maintained in an incubator (37°C, 5% CO₂) to preserve their structural integrity. To induce spheroid collapse, the structures were cooled to room temperature (≤25°C) for 5–10 minutes.

### ^51^Cr release assay

Cytotoxicity of NK cell lines YTS and NK92 was assessed using a ^51^Cr-release assay with target cell lines 721.221 and K562, respectively. Target cells were labeled with Na_2_^51^CrO_4_ (100 µCi per 10^6^ cells) at 37°C for 1 hour. After labeling, the cells were washed three times and resuspended in complete R10 medium at a concentration of 10^5^ cells/ml. A total of 10^4^ ^51^Cr-labeled target cells were added to each well of U-bottom 96-well plates (Corning), mixed with NK cells at varying effector-to-target (E/T) ratios in triplicate, and incubated at 37°C for 4 hours. To determine maximal ^51^Cr release, 1% IGEPAL (v/v, Sigma-Aldrich) was used to lyse all cells. Spontaneous release was measured by incubating the ^51^Cr-labeled target cells in medium alone, while experimental release was determined from target cells co-incubated with NK cells. After incubation, the plates were centrifuged, and 100 µl of supernatant was transferred to LumaPlate-96 plates (PerkinElmer), air-dried, and measured using a TopCount NXT detector(PerkinElmer). The percentage of specific lysis was calculated with the formula below:

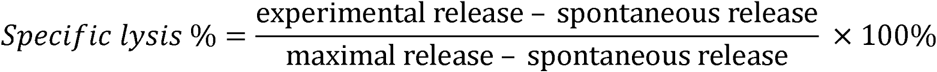

### Live Cell Imaging in 3D Models

To map the multidirectional killing activity of NK cells within the 3D model, both NK- triggering 721.221 cells and bystander K562 cells were labeled with 1 µM of the cytosolic dye Calcein Red-Orange AM (Invitrogen). Simultaneously, 721.221 cells were labeled with 1 µM CellMask Green (Invitrogen) and K562 cells with 1 µM CellMask Deep Red (Invitrogen) for membrane staining to distinguish between the two. All staining procedures were performed at 37°C for 30 minutes, followed by two washes with warm phenol red-free RPMI 1640 medium (Gibco). The labeled cells were then used to create cell strata and cultured for 4 hours following the temperature-triggered gel collapse. After incubation, the assembled gel was carefully cut open. The exposed sections were imaged using a Zeiss Axio Observer CSU-X spinning disc confocal microscope with a 100×/1.42 NA objective. Fluorescent signals were excited using 488 nm (CellMask Green), 561 nm (Calcein Red-Orange AM), and 640 nm (CellMask Deep Red) laser lines. Z-stack images were acquired with an interval of 0.4 µm over a depth of 30 µm to capture the entire 3D structure. Exposure time for each channel was set to 50∼100 ms. The acquired images were analyzed and quantified using FIJI (ImageJ, version 2.0 with default plugin).

## Key resources table

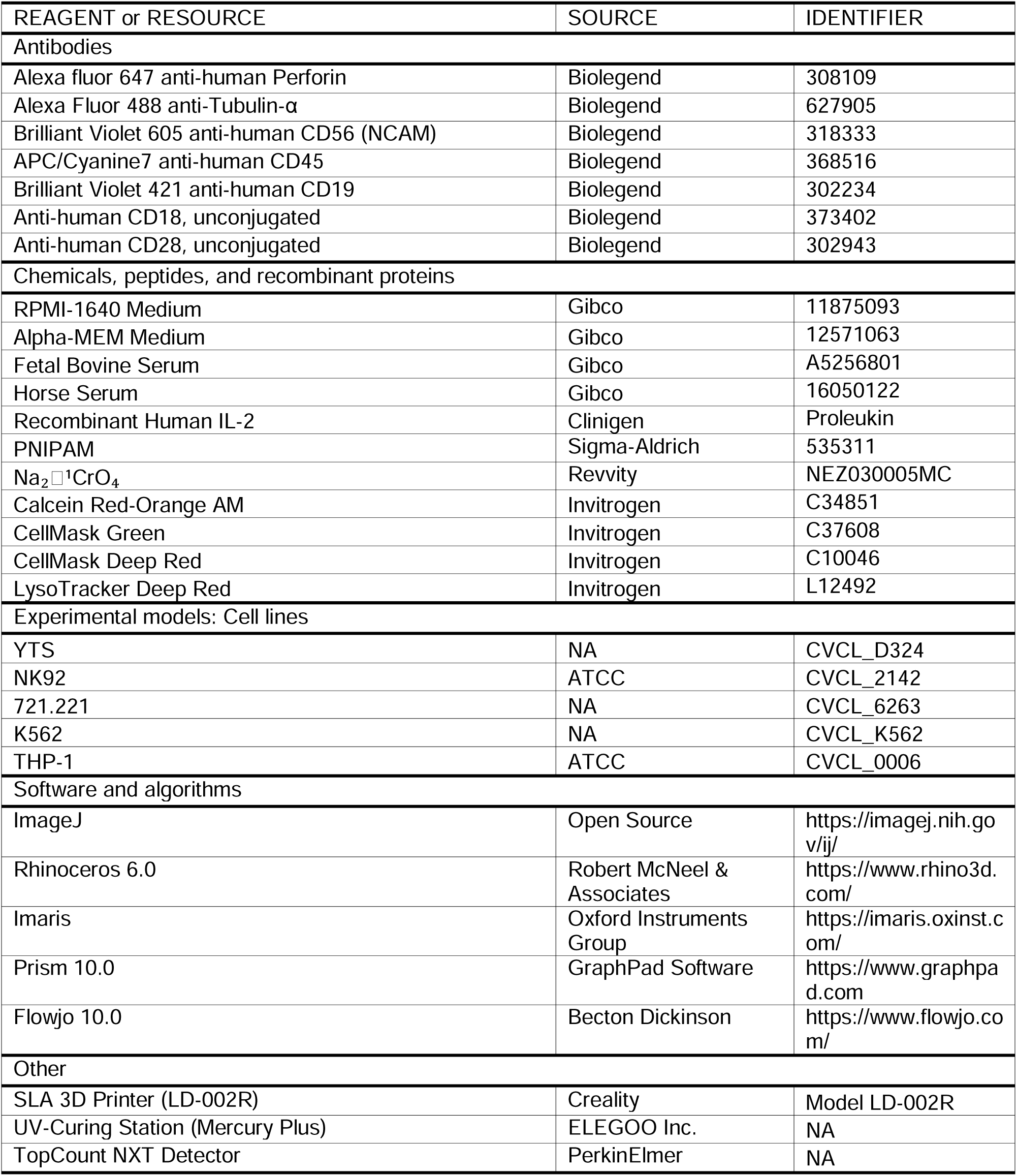

## References

1. Marofi F, Motavalli R, Safonov VA, Thangavelu L, Yumashev AV, Alexander M, et al. CAR T cells in solid tumors: challenges and opportunities. Stem Cell Res Ther. 2021;12(1):81.

2. Hui E. Immune checkpoint inhibitors. J Cell Biol. 2019;218(3):740–1.

3. Rodriguez-Garcia A, Palazon A, Noguera-Ortega E, Powell DJ, Guedan S. CAR-T Cells Hit the Tumor Microenvironment: Strategies to Overcome Tumor Escape. Front Immunol. 2020;11:1109.

4. Thistlethwaite FC, Gilham DE, Guest RD, Rothwell DG, Pillai M, Burt DJ, et al. The clinical efficacy of first-generation carcinoembryonic antigen (CEACAM5)-specific CAR T cells is limited by poor persistence and transient pre-conditioning-dependent respiratory toxicity. Cancer Immunol Immunother. 2017;66(11):1425–36.

5. Kershaw MH, Westwood JA, Parker LL, Wang G, Eshhar Z, Mavroukakis SA, et al. A phase I study on adoptive immunotherapy using gene-modified T cells for ovarian cancer. Clin Cancer Res. 2006;12(20 Pt 1):6106-15.

6. O’Rourke DM, Nasrallah MP, Desai A, Melenhorst JJ, Mansfield K, Morrissette JJD, et al. A single dose of peripherally infused EGFRvIII-directed CAR T cells mediates antigen loss and induces adaptive resistance in patients with recurrent glioblastoma. Sci Transl Med. 2017;9(399).

7. Albelda SM. CAR T cell therapy for patients with solid tumours: key lessons to learn and unlearn. Nat Rev Clin Oncol. 2024;21(1):47–66.

8. Fontana F, Marzagalli M, Sommariva M, Gagliano N, Limonta P. In Vitro 3D Cultures to Model the Tumor Microenvironment. Cancers (Basel). 2021;13(12).

9. Multhoff G, Vaupel P. Hypoxia Compromises Anti-Cancer Immune Responses. Adv Exp Med Biol. 2020;1232:131–43.

10. Jing X, Yang F, Shao C, Wei K, Xie M, Shen H, et al. Role of hypoxia in cancer therapy by regulating the tumor microenvironment. Mol Cancer. 2019;18(1):157.

11. Nia HT, Munn LL, Jain RK. Physical traits of cancer. Science. 2020;370(6516).

12. Guillaume L, Rigal L, Fehrenbach J, Severac C, Ducommun B, Lobjois V. Characterization of the physical properties of tumor-derived spheroids reveals critical insights for pre-clinical studies. Sci Rep. 2019;9(1):6597.

13. Cox TR. The matrix in cancer. Nat Rev Cancer. 2021;21(4):217–38.

14. Giussani M, Triulzi T, Sozzi G, Tagliabue E. Tumor Extracellular Matrix Remodeling: New Perspectives as a Circulating Tool in the Diagnosis and Prognosis of Solid Tumors. Cells. 2019;8(2).

15. Pathria P, Louis TL, Varner JA. Targeting Tumor-Associated Macrophages in Cancer. Trends Immunol. 2019;40(4):310–27.

16. Kazemi MH, Sadri M, Najafi A, Rahimi A, Baghernejadan Z, Khorramdelazad H, et al. Tumor- infiltrating lymphocytes for treatment of solid tumors: It takes two to tango? Front Immunol. 2022;13:1018962.

17. Guan X, Huang S. Advances in the application of 3D tumor models in precision oncology and drug screening. Front Bioeng Biotechnol. 2022;10:1021966.

18. Napoli GC, Figg WD, Chau CH. Functional Drug Screening in the Era of Precision Medicine. Front Med (Lausanne). 2022;9:912641.

19. Ramachandramoorthy H, Dang T, Srinivasa A, Nguyen KT, Nguyen P. Development of a Smart Portable Hypoxic Chamber with Accurate Sensing, Control and Visualization of In Vitro Cell Culture for Replication of Cancer Microenvironment. Cancers (Basel). 2023;15(14).

20. Ayuso JM, Rehman S, Farooqui M, Virumbrales-Muñoz M, Setaluri V, Skala MC, et al. Microfluidic Tumor-on-a-Chip Model to Study Tumor Metabolic Vulnerability. Int J Mol Sci. 2020;21(23).

21. Ligorio C, Mata A. Synthetic extracellular matrices with function-encoding peptides. Nat Rev Bioeng. 2023:1–19.

22. Xing H, Lee H, Luo L, Kyriakides TR. Extracellular matrix-derived biomaterials in engineering cell function. Biotechnol Adv. 2020;42:107421.

23. Teijeira A, Migueliz I, Garasa S, Karanikas V, Luri C, Cirella A, et al. Three-dimensional colon cancer organoids model the response to CEA-CD3 T-cell engagers. Theranostics. 2022;12(3):1373–87.

24. Gunti S, Hoke ATK, Vu KP, London NR. Organoid and Spheroid Tumor Models: Techniques and Applications. Cancers (Basel). 2021;13(4).

25. Gilazieva Z, Ponomarev A, Rutland C, Rizvanov A, Solovyeva V. Promising Applications of Tumor Spheroids and Organoids for Personalized Medicine. Cancers (Basel). 2020;12(10).

26. Sofia BL, Madalena ZDRFC, Donatella C, Sandra C, Marcelle H, Raffaella C, et al. Establishing the scientific validity of complex in vitro models. Luxembourg: Publications Office of the European Union; 2021.

27. Zhou Z, Pang Y, Ji J, He J, Liu T, Ouyang L, et al. Harnessing 3D in vitro systems to model immune responses to solid tumours: a step towards improving and creating personalized immunotherapies. Nat Rev Immunol. 2024;24(1):18–32.

28. Lugand L, Mestrallet G, Laboureur R, Dumont C, Bouhidel F, Djouadou M, et al. Methods for Establishing a Renal Cell Carcinoma Tumor Spheroid Model With Immune Infiltration for Immunotherapeutic Studies. Front Oncol. 2022;12:898732.

29. Sherman H, Gitschier HJ, Rossi AE. A Novel Three-Dimensional Immune Oncology Model for High-Throughput Testing of Tumoricidal Activity. Front Immunol. 2018;9:857.

30. Rodríguez CF, Andrade-Pérez V, Vargas MC, Mantilla-Orozco A, Osma JF, Reyes LH, et al. Breaking the clean room barrier: exploring low-cost alternatives for microfluidic devices. Front Bioeng Biotechnol. 2023;11:1176557.

31. Niculescu AG, Chircov C, Bîrcă AC, Grumezescu AM. Fabrication and Applications of Microfluidic Devices: A Review. Int J Mol Sci. 2021;22(4).

32. Fathi I, Imura T, Inagaki A, Nakamura Y, Nabawi A, Goto M. Decellularized Whole-Organ Pre-vascularization: A Novel Approach for Organogenesis. Front Bioeng Biotechnol. 2021;9:756755.

33. Giobbe GG, Crowley C, Luni C, Campinoti S, Khedr M, Kretzschmar K, et al. Extracellular matrix hydrogel derived from decellularized tissues enables endodermal organoid culture. Nat Commun. 2019;10(1):5658.

34. Cao H, Duan L, Zhang Y, Cao J, Zhang K. Current hydrogel advances in physicochemical and biological response-driven biomedical application diversity. Signal Transduct Target Ther. 2021;6(1):426.

35. Jeong B, Kim SW, Bae YH. Thermosensitive sol-gel reversible hydrogels. Adv Drug Deliv Rev. 2002;54(1):37–51.

36. Haq MA, Su Y, Wang D. Mechanical properties of PNIPAM based hydrogels: A review. Mater Sci Eng C Mater Biol Appl. 2017;70(Pt 1):842–55.

37. Carter JM, Polley MC, Leon-Ferre RA, Sinnwell J, Thompson KJ, Wang X, et al. Characteristics and Spatially Defined Immune (micro)landscapes of Early-stage PD-L1-positive Triple-negative Breast Cancer. Clin Cancer Res. 2021;27(20):5628–37.

38. Mentlik AN, Sanborn KB, Holzbaur EL, Orange JS. Rapid lytic granule convergence to the MTOC in natural killer cells is dependent on dynein but not cytolytic commitment. Mol Biol Cell. 2010;21(13):2241–56.

39. Hsu HT, Mace EM, Carisey AF, Viswanath DI, Christakou AE, Wiklund M, et al. NK cells converge lytic granules to promote cytotoxicity and prevent bystander killing. J Cell Biol. 2016;215(6):875–89.

40. McCann FE, Vanherberghen B, Eleme K, Carlin LM, Newsam RJ, Goulding D, et al. The size of the synaptic cleft and distinct distributions of filamentous actin, ezrin, CD43, and CD45 at activating and inhibitory human NK cell immune synapses. J Immunol. 2003;170(6):2862–70.

41. Dhamecha D, Le D, Chakravarty T, Perera K, Dutta A, Menon JU. Fabrication of PNIPAm-based thermoresponsive hydrogel microwell arrays for tumor spheroid formation. Mater Sci Eng C Mater Biol Appl. 2021;125:112100.

42. Dosh RH, Essa A, Jordan-Mahy N, Sammon C, Le Maitre CL. Use of hydrogel scaffolds to develop an in vitro 3D culture model of human intestinal epithelium. Acta Biomater. 2017;62:128–43.

43. Dosh RH, Jordan-Mahy N, Sammon C, Le Maitre CL. Use of l-pNIPAM hydrogel as a 3D-scaffold for intestinal crypts and stem cell tissue engineering. Biomater Sci. 2019;7(10):4310–24.

44. Badalamenti G, Fanale D, Incorvaia L, Barraco N, Listì A, Maragliano R, et al. Role of tumor-infiltrating lymphocytes in patients with solid tumors: Can a drop dig a stone? Cell Immunol. 2019;343:103753.

45. Boucherit N, Gorvel L, Olive D. 3D Tumor Models and Their Use for the Testing of Immunotherapies. Front Immunol. 2020;11:603640.

46. Bhat SM, Badiger VA, Vasishta S, Chakraborty J, Prasad S, Ghosh S, et al. 3D tumor angiogenesis models: recent advances and challenges. J Cancer Res Clin Oncol. 2021;147(12):3477–94.

47. Zurowski D, Patel S, Hui D, Ka M, Hernandez C, Love AC, et al. High-throughput method to analyze the cytotoxicity of CAR-T Cells in a 3D tumor spheroid model using image cytometry. SLAS Discov. 2023;28(3):65–72.

48. Grunewald L, Lam T, Andersch L, Klaus A, Schwiebert S, Winkler A, et al. A Reproducible Bioprinted 3D Tumor Model Serves as a Preselection Tool for CAR T Cell Therapy Optimization. Front Immunol. 2021;12:689697.

49. Song J, Choi H, Koh SK, Park D, Yu J, Kang H, et al. High-Throughput 3D. Front Immunol. 2021;12:733317.

50. Rodrigues J, Heinrich MA, Teixeira LM, Prakash J. 3D In Vitro Model (R)evolution: Unveiling Tumor-Stroma Interactions. Trends Cancer. 2021;7(3):249–64.

51. Vitale I, Manic G, Coussens LM, Kroemer G, Galluzzi L. Macrophages and Metabolism in the Tumor Microenvironment. Cell Metab. 2019;30(1):36–50.

52. Hinshaw DC, Shevde LA. The Tumor Microenvironment Innately Modulates Cancer Progression. Cancer Res. 2019;79(18):4557–66.

53. Habanjar O, Diab-Assaf M, Caldefie-Chezet F, Delort L. 3D Cell Culture Systems: Tumor Application, Advantages, and Disadvantages. Int J Mol Sci. 2021;22(22).

54. Lueckgen A, Garske DS, Ellinghaus A, Mooney DJ, Duda GN, Cipitria A. Enzymatically-degradable alginate hydrogels promote cell spreading and in vivo tissue infiltration. Biomaterials. 2019;217:119294.

55. Wang D, Duan J, Liu J, Yi H, Zhang Z, Song H, et al. Stimuli-Responsive Self-Degradable DNA Hydrogels: Design, Synthesis, and Applications. Adv Healthc Mater. 2023;12(16):e2203031.

56. Macková H, Hlídková H, Kaberova Z, Proks V, Kučka J, Patsula V, et al. Thiolated poly(2- hydroxyethyl methacrylate) hydrogels as a degradable biocompatible scaffold for tissue engineering. Mater Sci Eng C Mater Biol Appl. 2021;131:112500.

57. Rosenfeld A, Göckler T, Kuzina M, Reischl M, Schepers U, Levkin PA. Designing Inherently Photodegradable Cell-Adhesive Hydrogels for 3D Cell Culture. Adv Healthc Mater. 2021;10(16):e2100632.

58. Villiou M, Paez JI, Del Campo A. Photodegradable Hydrogels for Cell Encapsulation and Tissue Adhesion. ACS Appl Mater Interfaces. 2020;12(34):37862–72.

59. Norris SCP, Soto J, Kasko AM, Li S. Photodegradable Polyacrylamide Gels for Dynamic Control of Cell Functions. ACS Appl Mater Interfaces. 2021;13(5):5929–44.

60. Raman R, Hua T, Gwynne D, Collins J, Tamang S, Zhou J, et al. Light-degradable hydrogels as dynamic triggers for gastrointestinal applications. Sci Adv. 2020;6(3):eaay0065.

61. Mehta P, Rahman Z, Ten Dijke P, Boukany PE. Microfluidics meets 3D cancer cell migration. Trends Cancer. 2022;8(8):683–97.

62. Steinman JB, Santarossa CC, Miller RM, Yu LS, Serpinskaya AS, Furukawa H, et al. Chemical structure-guided design of dynapyrazoles, cell-permeable dynein inhibitors with a unique mode of action. Elife. 2017;6.

63. Stinchcombe JC, Bossi G, Booth S, Griffiths GM. The immunological synapse of CTL contains a secretory domain and membrane bridges. Immunity. 2001;15(5):751–61.

64. Carisey AF, Mace EM, Saeed MB, Davis DM, Orange JS. Nanoscale Dynamism of Actin Enables Secretory Function in Cytolytic Cells. Curr Biol. 2018;28(4):489–502.e9.

65. Wang MS, Hu Y, Sanchez EE, Xie X, Roy NH, de Jesus M, et al. Mechanically active integrins target lytic secretion at the immune synapse to facilitate cellular cytotoxicity. Nat Commun. 2022;13(1):3222.

