## Supplementary material for "A Thermo-responsive collapse system for controlling heterogeneous cell localization, ratio and interaction for three-dimensional solid tumor modeling": Key Resources Table

| REAGENT or RESOURCE | SOURCE | IDENTIFIER |
| --- | --- | --- |
| Antibodies | | |
| Alexa fluor 647 anti-human Perforin | Biolegend | 308109 |
| Alexa Fluor 488 anti-Tubulin-α | Biolegend | 627905 |
| Brilliant Violet 605 anti-human CD56 (NCAM) | Biolegend | 318333 |
| APC/Cyanine7 anti-human CD45 | Biolegend | 368516 |
| Brilliant Violet 421 anti-human CD19 | Biolegend | 302234 |
| Anti-human CD18, unconjugated | Biolegend | 373402 |
| Anti-human CD28, unconjugated | Biolegend | 302943 |
| Chemicals, peptides, and recombinant proteins | | |
| RPMI-1640 Medium | Gibco | 11875093 |
| Alpha-MEM Medium | Gibco | 12571063 |
| Fetal Bovine Serum | Gibco | A5256801 |
| Horse Serum | Gibco | 16050122 |
| Recombinant Human IL-2 | Clinigen | Proleukin |
| PNIPAM | Sigma-Aldrich | 535311 |
| Na₂⁵¹CrO₄ | Revvity | NEZ030005MC |
| Calcein Red-Orange AM | Invitrogen | C34851 |
| CellMask Green | Invitrogen | C37608 |
| CellMask Deep Red | Invitrogen | C10046 |
| LysoTracker Deep Red | Invitrogen | L12492 |
| Experimental models: Cell lines | | |
| YTS | NA | CVCL_D324 |
| NK92 | ATCC | CVCL_2142 |
| 721.221 | NA | CVCL_6263 |
| K562 | NA | CVCL_K562 |
| THP-1 | ATCC | CVCL_0006 |
| Software and algorithms | | |
| ImageJ | Open Source | https://imagej.nih.gov/ij/ |
| Rhinoceros 6.0 | Robert McNeel & Associates | https://www.rhino3d.com/ |
| Imaris | Oxford Instruments Group | https://imaris.oxinst.com/ |
| Prism 10.0 | GraphPad Software | https://www.graphpad.com |
| Flowjo 10.0 | Becton Dickinson | https://www.flowjo.com/ |
| Other | | |
| SLA 3D Printer (LD-002R) | Creality | Model LD-002R |
| UV-Curing Station (Mercury Plus) | ELEGOO Inc. | NA |
| TopCount NXT Detector | PerkinElmer | NA |
